# TGF-β broadly modifies rather than specifically suppresses reactivated memory CD8 T cells in a dose-dependent manner

**DOI:** 10.1101/2023.07.27.550871

**Authors:** Alexis Taber, Andrew Konecny, James Scott-Browne, Martin Prlic

## Abstract

Transforming growth factor β (TGF-β) directly acts on naïve, effector and memory T cells to control cell fate decisions, which was shown using genetic abrogation of TGF-β signaling. TGF-β availability is altered by infections and cancer, however the dose-dependent effects of TGF-β on memory CD8 T cell (T_mem_) reactivation are still poorly defined. We examined how activation and TGF-β signals interact to shape the functional outcome of T_mem_ reactivation. We found that TGF-β could suppress cytotoxicity in a manner that was inversely proportional to the strength of the activating TCR or pro-inflammatory signals. In contrast, even high doses of TGF-β had a comparatively modest effect on IFN-γ expression in the context of weak and strong reactivation signals. Since CD8 T_mem_ may not always receive TGF-β signals concurrently with reactivation, we also explored whether the temporal order of reactivation versus TGF-β signals is of importance. We found that exposure to TGF-β prior to as well as after an activation event were both sufficient to reduce cytotoxic effector function. Concurrent ATAC-seq and RNA-seq analysis revealed that TGF-β altered ∼10% of the regulatory elements induced by reactivation and also elicited transcriptional changes indicative of broadly modulated functional properties. We confirmed some changes on the protein level and found that TGF-β-induced expression of CCR8 was inversely proportional to the strength of the reactivating TCR signal. Together, our data suggest that TGF-β is not simply suppressing CD8 T_mem_, but modifies functional and chemotactic properties in context of their reactivation signals and in a dose-dependent manner.

## Introduction

The pleiotropic functions of TGF-β have been described in a wealth of literature and include roles in angiogenesis, wound healing, cancer and regulating immune responses (1, 2). TGF-β1 (often referred to as TGF-β since it is the most prevalent and studied isoform (3)) is typically considered to be a powerful suppressor of the immune response (3). Most immune cells express the TGFβ type I and type II serine/threonine kinase receptors (also referred to as TβRI and TβR2, or TGF-βRI and TGF-βRII) and are thus able to respond to TGF-β signals (2). TGF-β affects T cells at all stages of development, starting in the thymus during T cell development, T cell homeostasis in the periphery, as well as T cell differentiation following activation (4). Two genetic models have been widely used to define the consequences of TGF-β signaling on T cell fate and differentiation: First, transgenic mice expressing a dominant negative form of the TGF-β receptor II (dnTGβRII) under the control of the CD4 promoter that lacks the CD8 silencer (5) or a CD2 promoter (6), thus allowing for expression in CD4 and CD8 T cells in these mouse lines. A follow up study with the CD4-dnTGFβRII mice revealed that the dominant negative receptor still had some signaling capacity (possibly independent of bona fide TGF-β receptor activation) (7), which somewhat complicates interpretation of studies that used these mice. Second, mice bearing *TGF-βr2* alleles with flanking loxP sites (floxed TGF-βRII) allow for conditional deletion in T cells by crossing to mice expressing Cre recombinase under control of the *Cd4* promoter that is active in thymocytes (CD4-cre) (8, 9), or expressing Cre under control of the distal Lck promoter active in mature, naïve T cells (dLck-cre) (10). Of note, these distinct approaches to abrogating TGF-β signaling in T cells also had distinct disease phenotypes, which ultimately helped separate the roles of TGF-β signals during thymic selection and maintenance of tolerance in the periphery (8-10). To study the consequences of TGF-β signals during the effector stage flox TGF-βRII mice were crossed to mice expressing Cre under control of the Granzyme B locus (granzyme B-cre), which revealed a role for TGF-β in controlling the number of short lived effector cells (7). To study the effect on T_mem_, flox TGF-βRII mice were crossed to mice expressing Cre fused to the ligand binding domain of the estrogen receptor (ER-cre) (11). Tamoxifen-induced Cre mediated deletion of TGF-βRII during the CD8 T_mem_ stage revealed that TGF-β signals are required for the maintenance of the CD8 T_mem_ transcriptional program and function (11).

An inherent limitation of these powerful genetic approaches is that deletion or expression of a dominant negative form of a receptor precludes studying dose-dependent effects of the ligand. For cytokines and T cells, it is noteworthy that the effect of a signal on T cell fate decisions does not necessarily follow a titration curve, but can result in a quantal – all or none – outcome (12). In context of TGF-β, the potential dose-dependent effects on T cells at various stages of differentiation are still poorly defined. This is at least in part due to the challenge of measuring biologically active TGF-β (13). TGF-β is abundant in blood and tissues, but most of the TGF-β in blood and tissues is present as a complex with latency associated peptide (LAP) and latent TGF-β-binding proteins (LTBPs), respectively. Once activated by integrins or other signals, the receptor-binding site of TGF-β is exposed and TGF-β becomes active. Measuring the availability of the latent and active form of TGF-β is possible using ELISAs and reporter cells (13-15), but often varies based on the reagents and protocols used (13). Thus, the concentration range of biologically active TGF-β in health versus disease is still poorly defined. We were specifically interested in potential concentration-dependent effects of TGF-β on CD8 T_mem_ in the context of reactivation. CD8 T_mem_ reactivation is typically considered in the context of repeated infections with pathogens, but is also highly relevant in the context of tumor responses: vaccines that elicit immune responses against tumor antigens have had promising results and generate memory T cells (16, 17), and tumor-specific memory CD8 T cells responding to PD-1/PD-L1 blockade reside in the tumor draining lymph node(18). We thus wanted to examine if the effect of TGF-β on memory CD8 T cell reactivation is (1) potentially distinct from its role during T cell priming, and (2) TGF-β dose- and (3) activation-signal dependent. Since TGF-β has been reported to inhibit IFN-γ production by cytokine-activated memory CD8 T cells (19, 20), we wanted to define if the type of activating signal (T cell receptor- vs. cytokine-mediated) yields distinct responses to TGF-β signals.

Since the mouse model has been so widely used to define the effects of TGF-β signaling, we also used a mouse model system to generate a CD8 T_mem_ population with expressing a well-defined T cell receptor specific for an epitope of chicken ovalbumin (OT-I T cells). We utilized OT-I T_mem_ to define how low to high concentrations of TGF-β signals affect the functional properties of CD8 T_mem_ across a range of reactivation signals (weak to strong TCR activating, and cytokine-driven activation). We found that TGF-β was not broadly immunosuppressive, but rather altered functional and chemotactic properties in a dose- and reactivation context-dependent manner. TGF-β could suppress cytotoxicity in a manner that was inversely proportional to the strength of the activating TCR or pro-inflammatory signal. In contrast, TGF-β had a rather modest effect on IFN-γ expression. Importantly, TGF-β was not merely suppressing aspects of effector function, it directly increased expression of some chemokine receptors, including CCR8. TGF-β induced the expression of CCR8 in CD8 T_mem_ regardless if reactivation occurred via TCR or cytokines. Interestingly, induction of expression was inversely proportional to the suppression in cytotoxicity and most effective in CD8 T_mem_ reactivated by a weak TCR signal. We discuss the implication of our findings in context of CD8 T_mem_ reactivation in response to infections and in tumors.

## Results

### TGF-β strongly inhibits cytotoxic function, but not IFN-γ production by CD8 T_mem_ in a dose-dependent manner

First, we sought to determine whether TGF-β affected the function of reactivated CD8 T_mem_. To generate a population of CD8 T_mem_ with known Ag-specificity, we transferred congenically marked OT-I T cells, which recognize the SIINFEKL (N4) epitope of chicken ovalbumin (OVA) bound to the MHC class I molecule H-2K^b^, into C57BL/6J mice. We then infected these mice with OVA-expressing vesicular stomatitis virus (VSV-OVA) and waited at least 60 days before isolating cells from these OT-I memory mice. As a first step, we wanted to define the effect of a high dose (100 ng/ml) of TGF-β in context of a very strong reactivating signal: we isolated T cells from spleen and lymph nodes (LN) of OT-I memory mice followed by ex vivo stimulation with plate-bound anti-CD3/28 antibodies (CD3/28) for 24 hours with or without TGF-β (**Figure 1A**). Using flow cytometry, we found that TGF-β was sufficient to reduce IFN-γ expression, but greatly diminished GzmB expression in reactivated OT-I memory T cells (**Figure 1B and 1C**). Next, we titrated the concentration of TGF-β to assess its dose-dependent effects. As a reference value, the total TGF-β1 in mouse spleen has been reported to be ∼1000ng/g spleen (21) (**Supplemental Figure 1A**), but active TGF-β is often only a fraction of total TGF-β (13). We observed again that IFN-γ was fairly resistant to TGF-β as only concentrations above 1ng/ml appeared to have at least a modest effect (**Figure 1D**). In contrast, cytotoxicity was much more susceptible to TGF-β-mediated suppression as a dose of 1.3ng/ml was sufficient to decrease the frequency of granzyme B expressing OT-I T cells 2-fold, indicated as the “1/2 Max” value. We also examined activation associated protein biomarkers in OT-I T_mem_ and found that frequency of cells expressing Programmed Death 1 (PD-1) and median fluorescence intensity (MedFI) of the transcription factor T cell factor 1 (TCF1) were very modestly but significantly increased by TGF-β. In contrast, the frequency of Ki67 expressing T_mem_ and the MedFI of TOX did not significantly change (**Supplemental Figure 1B**). While PD-1 is characteristically a target for T cell inhibition, its upregulation does not necessarily connote T cell “exhaustion” (22).

**Figure 1.**
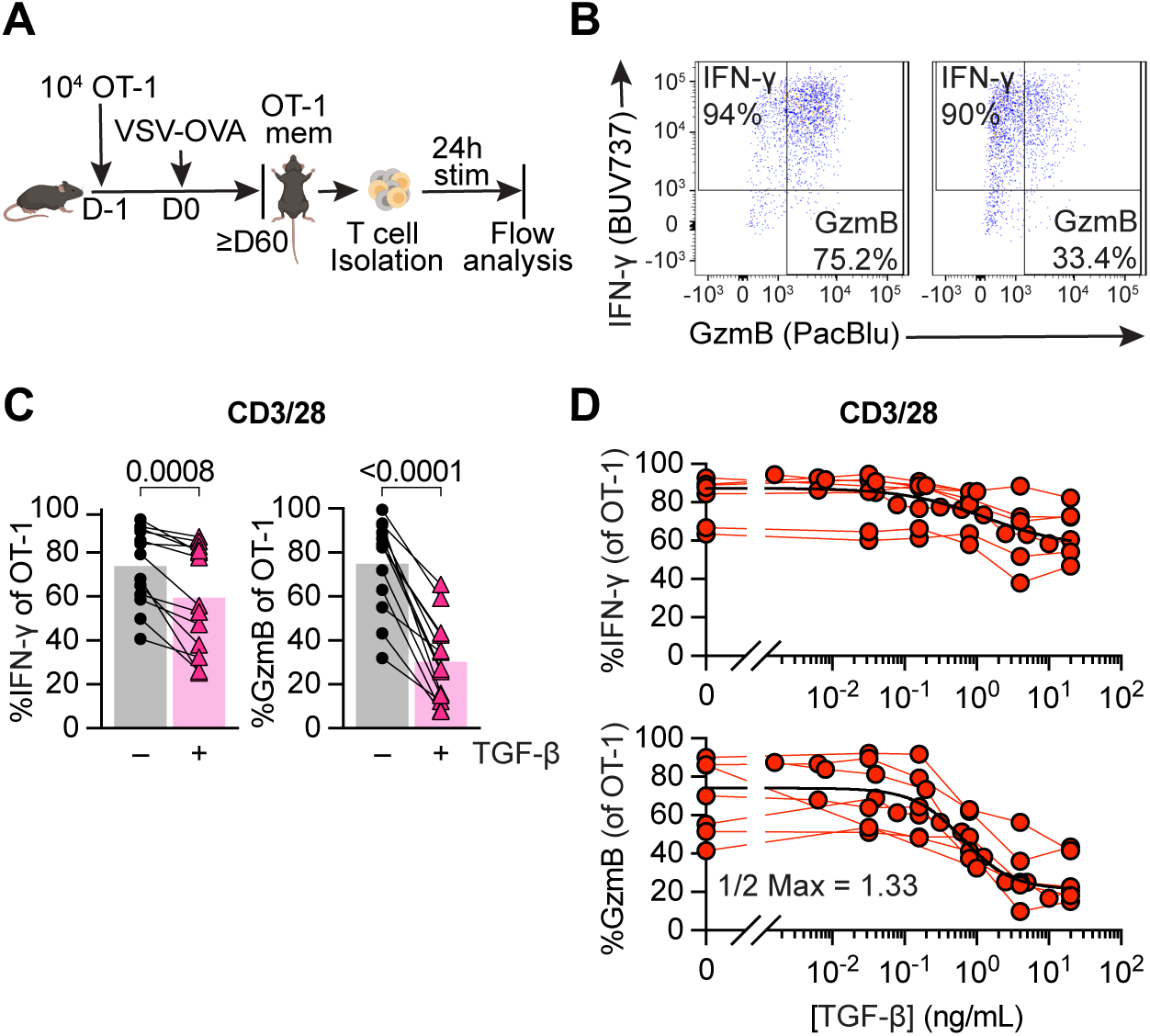
TGF-β preferentially inhibits cytotoxicity of memory CD8+ T cells in a dose-dependent manner. **(A**) Schematic of naïve OT-I CD8+ T cell adoptive transfer, memory OT-I T cell generation with VSV-OVA, T cell isolation with magnet-activated cell sorting (MACS) from Ag-experienced OT-I memory mice, and subsequent ex vivo stimulation and analysis. Stimulation was 24 hours with plate-bound anti-CD3 and anti-CD28 (CD3/28) in the presence or absence of TGF-β at 100ng/mL. **(B)** Representative expression and gating of IFN-γ and GzmB in OT-I T_mem_ post stimulation. **(C**) IFN-γ and GzmB frequencies. Each point represents an individual animal, with connecting lines across points from the same animal (n = 14 animals). Statistical significances were calculated using paired t tests. **(D)** Frequencies of IFN-γ and GzmB in OT-I T_mem_ post 24 hours stimulation with CD3/28 in the presence of titrated TGF-β (n = 7). TGF-β was titrated in two-fold dilutions starting with 20ng/mL and ending with 0.04ng/mL, in five-fold dilutions starting with 20ng/mL and ending at 0.032ng/mL, and in five-fold dilutions starting with 1ng/mL and ending at 0.0016ng/mL. Each point represents an individual animal with connecting lines across points from the same animal. Data shown are from 6 to 14 independent experiments.

Finally, we also assessed the effect on reactivation of endogenous CD8 T_mem_. We found that in the presence of TGF-β, reactivated endogenous T_mem_ had modestly reduced IFN-γ, but starkly decreased GzmB frequencies (**Supplemental Figure 1C**) thus mirroring our OT-I T cell data. Similarly, we found that the frequency of PD-1+ CD8 T_mem_ slightly increased, while frequency of Ki67, MedFI Tox, and MedFI TCF1 did not change (**Supplemental Figure 1D**). We observed similar effects of TGF-β when we recapitulated our ex vivo experimental approach with human CD8 T_mem_ (**Supplemental Figure 2A**). Of note, human PBMC contain effector memory CD8 T cells which express granzyme B prior to reactivation. TGF-β did not appear to affect this steady state expression pattern (**Supplemental Figure 2B**).

### TGF-β is not sufficient to fully suppress cytotoxicity when CD8+ T_mem_ are reactivated by strong TCR signals or cytokines

Cross-linking of the TCR by a monoclonal antibody delivers a very strong reactivation signal. To assess the effects of TGF-β in cells reactivated via their TCR triggered through peptide/MHC complexes, we compared OT-I T_mem_ reactivated by SIINFEKL (N4) and SIIQFEKL (Q4) peptides. N4 (SIINFEKL) bound to H-2K^b^ is a strong agonist for OT-I T cells, while the variant Q4 (SIIQFEKL) binds equally well to H-2K^b^ but is only a weak agonist for OT-I T cells (23). As an alternative reactivation signal, we also stimulated OT-I T_mem_ with a combination of IL-12, IL-15, and IL-18 (IL-12/15/18; Cyt) to induce reactivation in a TCR agonist-independent manner.

We found that IFN-γ expression was again only modestly affected in all experimental conditions (**Figure 2A**), while TGF-β essentially ablated cytotoxic function in N4- and Q4-reactivated OT-I T_mem_ (**Figure 2A; Suppl Figure 1E and 1F**). Cytokine-mediated reactivation yielded outcomes comparable to TCR cross-linking with and without cytokine treatment (**Suppl Figure 1E and 1F**). Finally, we titrated TGF-β in context of reactivation with N4, Q4, and Cyt stimulation (**Figure 2B and 2C**). When we reactivated OT-I T_mem_ with either N4 or Q4 peptide, we found that a much lower concentration of TGF-β was sufficient to reduce granzyme B expression (0.16 and 0.09 ng/ml of TGF-β reduce the frequency of gzmB+ OT-I T_mem_ 2x fold for N4 and Q4, respectively compared to 0.99 ng/ml after cytokine reactivation), but the impact on IFN-γ expression was again much more limited across all restimulation conditions.

**Figure 2.**
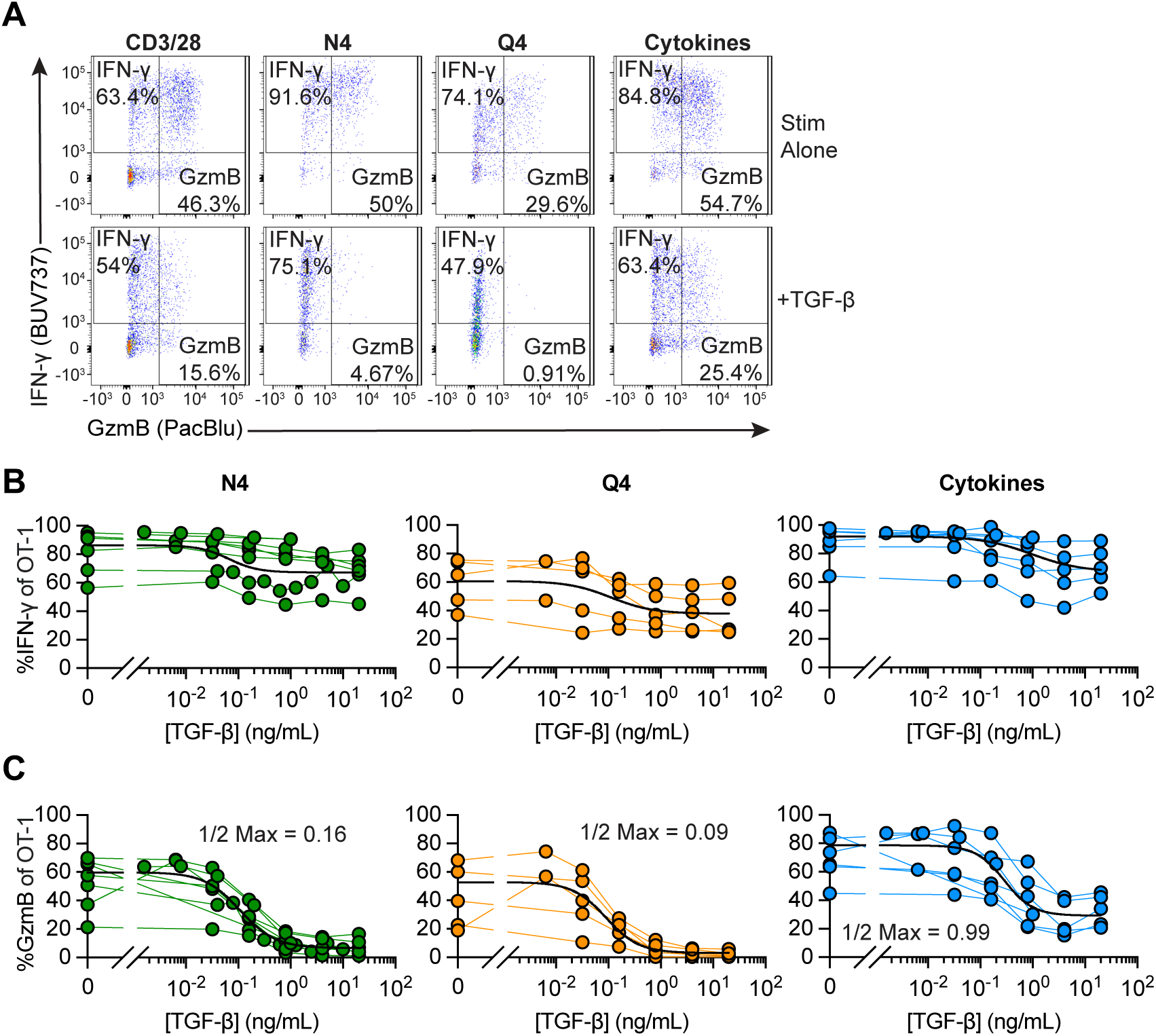
Strongly activated memory CD8+ T cells are less susceptible to TGF-β-mediated suppression. Stimulations were 24 hours with CD3/28, 100nM SIINFEKL (N4), 100nM SIIQFEKL (Q4), rIL-12, rIL-15, and rIL-18 in combination, with rIL12 and rIL-15 at 100ng/mL and rIL-18 at 0.5ng/mL (“Cyt”), and TGF-β at 20ng/mL. **(A)** Representative gating of IFN-γ and GzmB staining in OT-I T_mem_ at 24 hours post stimulation with CD3/28, N4, Q4, or Cyt in the presence or absence of TGF-β. **(B)** Frequencies of IFN-γ and **(C)** GzmB of OT-I T cells 24 hours post indicated stimulation condition in the presence of titrated TGF-β (N4 conditions are depicted from n = 7 animals, Q4 from n = 5, and Cytokines from n = 6). Each point represents an individual animal with connecting lines across points from the same animal. TGF-β was titrated in two-fold dilutions starting with 20ng/mL and ending with 0.04ng/mL, in five-fold dilutions starting with 20ng/mL and ending at 0.032ng/mL, and in five-fold dilutions starting with 1ng/mL and ending at 0.0016ng/mL (for N4 and Cyt). Calculated ½ Max inhibitory capacity values indicated. Data shown are from 4 to 6 independent experiments.

Finally, we also titrated the pro-inflammatory cytokines to determine the relationship between TGF-β and strength of the reactivating proinflammatory signals. We used IL-12/15/18 to elicit strong IFN-γ production and found that TGF-β could reduce IFN-γ 2-fold when less than 10ng/ml of each cytokine were available (**Suppl Fig 3**). To elicit strong granzyme B expression upon reactivation, we exposed OT-I T_mem_ to IL-12/IL-15 – granzyme B expression was again much more susceptible to inhibition by TGF-β and essentially completely inhibited unless more than 25ng/ml of each cytokine were present (**Suppl Fig 3, right**).

Together, these data indicate that the inhibitory effect of TGF-β on reactivation-induced cytotoxicity can be tuned by the concentration of TGF-β as well as the strength of the activating signal, while IFN-γ production is comparatively resistant to TGF-β-mediated suppression.

### TGF-β can still affect function if reactivation signals temporally precede the TGF-β signal

In these previous experiments we provided reactivation and TGF-β signals at the same time, but we considered that CD8 T_mem_ may receive activating signals before or after a TGF-β signal (for example, reactivation in a lymph node followed by a high dose TGF-β exposure in the tissue).

To test whether TGF-β could inhibit the cytotoxicity of already reactivated OT-I T_mem_, we modified the ex vivo stimulation conditions to include two additional experimental conditions: first reactivate the OT-I T_mem_, and then followed by adding TGF-β either 6 hours or 12 hours after the reactivation stimulation (TGF-β 6hrs, 12hrs; **Figure 3A**). We found that both the 6-hour and 12-hour delay between reactivation signal and TGF-β exposure inhibited IFN-γ to the same extent as the positive control (TGF at 0hrs) in N4- and Q4-activated memory CD8+ T cells (**Figure 3B**). IFN-γ was also attenuated by TGF-β when added 12 hours after anti-CD3/CD28-mediated reactivation, while a 12 hour delay had essentially no effect on IFN-γ production in the cytokine-mediated reactivation condition (**Figure 3B**). For GzmB, we found that the frequency of gzmB+ OT-I T_mem_ increased the longer the delay between reactivation and TGF-β addition for N4 and Q4-mediated reactivation. For the CD3/CD28 and cytokine stimulation conditions the 0hr control and 6hr delay groups were similar (**Figure 3C**), while OT-I T_mem_ in the 12 hour delay condition had more gzmB+ cells than the 0hr control group, but less than the no TGF-β control (**Figure 3C**). To determine if IFN-γ expression at the 24 hour analysis time point reflects an overall decrease in IFN-γ production or altered IFN-γ production kinetics, we measured the concentrations of IFN-γ in the culture supernatant. We observed similar trends of reduction in these experimental groups, but these were not statistically significant (**Supplemental Figure 4a**). Together, these data indicate that TGF-β can effectively limit cytotoxic function after CD8 T_mem_ have already been reactivated, particularly in context of reactivation with a low affinity ligand, while only modestly affecting IFN-γ production.

**Figure 3.**
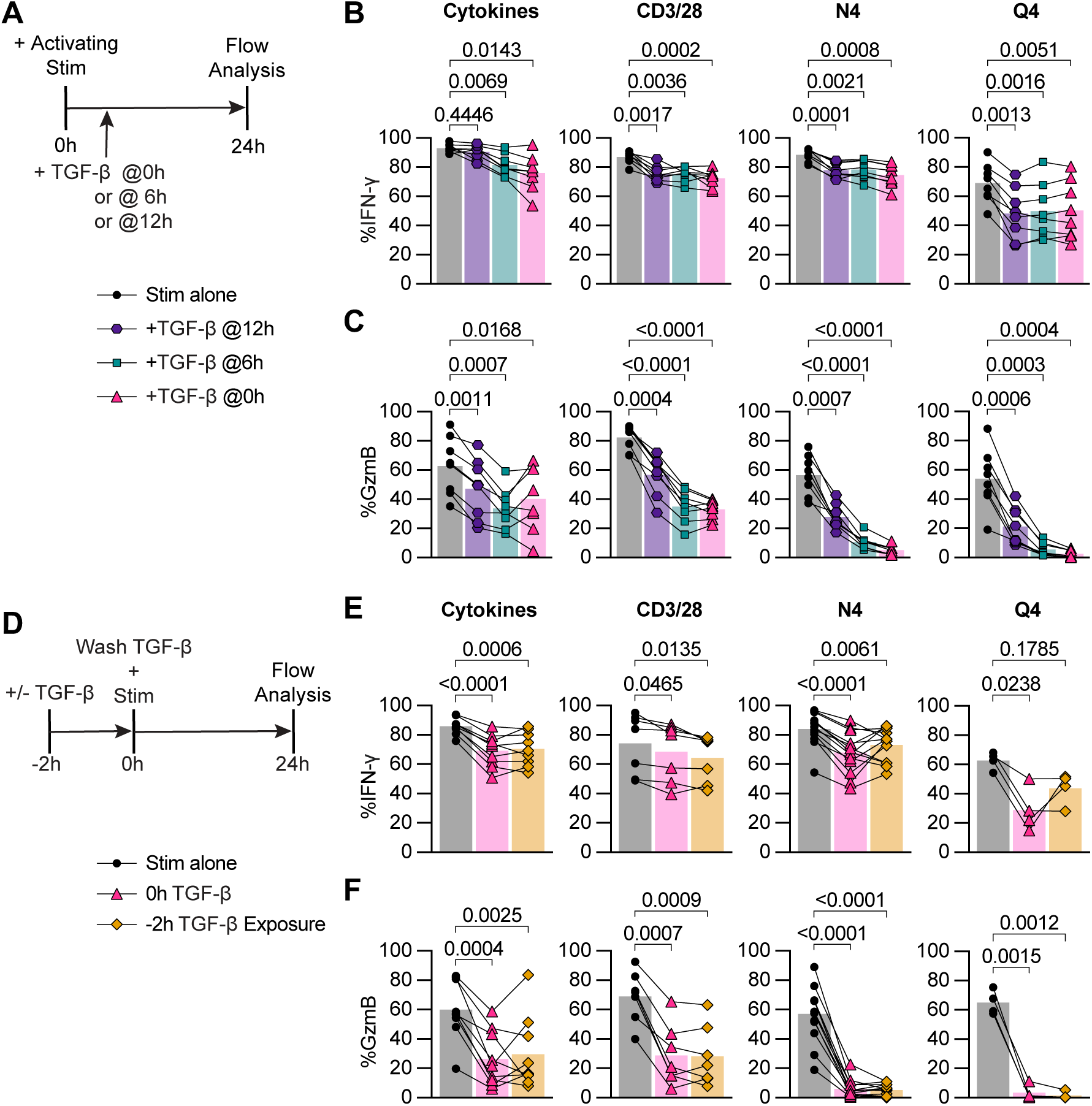
TGF-β inhibits cytotoxicity from recently reactivated memory CD8+ T cells and short-term exposure to TGF-β inhibits cytotoxicity of subsequently activated memory CD8+ T cells. **(A)** Schematic of ex vivo stimulation; cells were stimulated for 24 hours with CD3/28, N4, Q4, Cyt, and TGF-β at 100ng/mL. TGF-β was added 0 hours, 6 hours, or 12 hours post-start of activating stimulation. **(B)** Frequencies of IFN-γ and **(C)** GzmB in OT-I T_mem_ compared across stimulation conditions with TGF-β addition at indicated timepoints (n = 8 animals). **(D)** Schematic of ex vivo stimulation of isolated T cells from OT-I memory mice. Cells were treated with 100ng/mL TGF-β or media alone for 2 hours, the TGF-β was then washed out (down to 0.001ng/mL), immediately followed by 24 hours of activating stimulation. Stimulations were CD3/28, N4, Q4, Cyt, and TGF-β at 100ng/mL. **(E)** Frequencies of IFN-γ and **(F)** GzmB in OT-I T cells compared across stimulation conditions. CD3/28 data depicted are from n = 7 animals, N4 from n = 13, Q4 from n = 4, and Cyt from n = 9. All indicated statistical significances were calculated using one-way ANOVA with Tukey’s multiple comparison test. Data shown are from 3 to 10 independent experiments.

### Short-term exposure to TGF-β is sufficient to inhibit cytotoxicity of subsequently activated memory CD8+ T cells

Next, we reversed the order of signals and asked whether a brief exposure to TGF-β prior to reactivation could also inhibit the subsequent CD8 T_mem_ effector response (for example, exposure to TGF-β in the tissue prior to tissue egress into the draining LN). We pre-exposed OT-I T_mem_ to TGF-β for 2 hours, washed out the TGF-β, then stimulated these cells for 24 hours (**Figure 3D**). Interestingly, we found that IFN-γ was inhibited to a similar extent in the 2-hour pre-exposure condition as in the 0h (TGF-β added with stimulation for 24hrs) positive control TGF-β condition following cytokine stimulation (**Figure 3E**). Similarly, in the CD3/28 stimulation condition, an average of 74.3% of OT-I T_mem_ expressed IFN-γ, which decreased to 68.6% and 64.5% in the 0h TGF-β and 2h exposure conditions respectively. In the N4 and Q4 conditions, the pre-exposure had a different effect: IFN-γ expression increased from 67.9% to 73.1% (N4) and 29.0% to 43.8% (Q4) in the 0h TGF-β versus 2h pre-exposure conditions (**Figure 3E**). Surprisingly, we found that the 2h pre-exposure to TGF-β was sufficient to inhibit GzmB expression to the same drastic extent as prolonged TGF-β exposure. In the CD3/28 stimulation, GzmB expression, on average, decreased 2-fold from 69.07% to 28.72%, and 28.03% in the 0h TGF-β and 2h exposure conditions respectively. The frequency of granzyme B+ OT-I T_mem_ reactivated by N4 decreased from 57.04% to 6% with 0h TGF-β or 5.18% with 2h exposure. OT-I T_mem_ cells reactivated by Q4 had an even greater 20-fold reduction in GzmB expression, from 64.95% to 3.49% and 1.5% in the 0h TGF-β and 2h exposure conditions, respectively. In line with previous data, the cytokine activated OT-I T_mem_ exhibited suppression similar to CD8 T_mem_ reactivated by TCR cross-linking, with GzmB expression decreasing from 60.11% to only 26.45% and 29.60% in the 0h TGF-β and 2h exposure conditions. (**Figure 3F**).

We next examined whether the TGF-β exposed T_mem_ could regain full cytotoxic function after a short rest period. To test this, after the 2-hour exposure to TGF-β, we rested the T cells in fresh media for 4 hours before addition of activating stimulation for 24 hours. We found that even after resting for 4 hours post-TGF-β exposure GzmB expression could not be rescued, maintaining the 2-fold reduction in the CD3/28 condition, the 10-fold reduction in the N4 condition, and 20-fold reduction the Q4 condition (**Suppl Fig 4B**). In contrast, IFN-γ expression was similarly reduced when compared to the 0h and 2h exposure conditions (**Supp Fig 4C**). Overall, these data indicate that a short exposure to TGF-β is sufficient to control CD8 T_mem_ cytotoxic effector function for at least 24hrs. To elucidate how this may occur, we next examined how TGF-β alters chromatin accessibility and the transcriptome of CD8 T_mem_.

### Brief exposure to TGF-β is sufficient to epigenetically and transcriptionally alter memory CD8+ T cells

We next concomitantly interrogated the epigenetic and transcriptional effects of TGF-β on reactivated memory CD8+ T cells. We set up a short TGF-β exposure condition (+/- 2hrs of TGF-β in the absence of stimulation) and a 24hrs ex vivo restimulation condition (N4 +/- TGF-β). OT-I T_mem_ from the same experiment were analyzed in parallel using ATAC- and RNA-sequencing (**Figure 4A**). Of note, we also assessed granzyme B and IFN-γ protein expression in parallel, thus allowing us to link these protein, transcript, and epigenetic datasets. The ATAC-seq data revealed that a 2-hour exposure to TGF-β was sufficient to detect increases in chromatin accessibility at regions containing motifs bound by the SMAD family, which are the downstream transcriptional factors of the TGF-βR complex (24), compared to media alone (**Figure 4B**). The effect of TGF-β on SMAD TF motif associated chromatin accessibility was more pronounced - in the 24hrs stimulation condition (**Figure 4B**) and was accompanied by change in chromatin accessibility at more than 2500 regulatory elements (**Figure 4C**). Of note, reactivation itself altered nearly 26000 regulatory elements, which is about 25% of the regulatory elements in our global peak set of 99,317 peaks. Thus, TGF-β affects about 10% of the regulatory elements that are altered during reactivation. Similarly to the ATAC-seq data, we also detected some transcriptional changes in our +/- 2hr TGF-β group (28 down and 46 up), and a more substantial change (134 down 378 up) in the transcriptome after 24hrs of N4 stimulation +/- TGF-β (**Figure 4D and 4E**). Consistent with our flow cytometry findings, we found that GzmB was significantly decreased while IFN-γ was only minimally affected in the N4 + TGF-β condition at 24 hours (**Figure 4F**). Interestingly, we also found significant decreases in GzmC and Prf1 (**Figure 4F**). The ATAC-seq data indicate that there are no significant changes in accessibility for at the *Ifng*, *Gzmb*, *Gzmc* or *Prf1* loci, all of which have decreased transcriptional abundance in the N4 + TGF-β group (**Figure 4G**). As in our previous experiments, gzmB protein expression was decreased in these experiments as well **(Suppl Fig 5A.)**. Together, these data indicate that TGF-β can alter over 2500 regulatory elements during reactivation, but not all transcriptional changes are necessarily caused by epigenetic changes.

**Figure 4.**
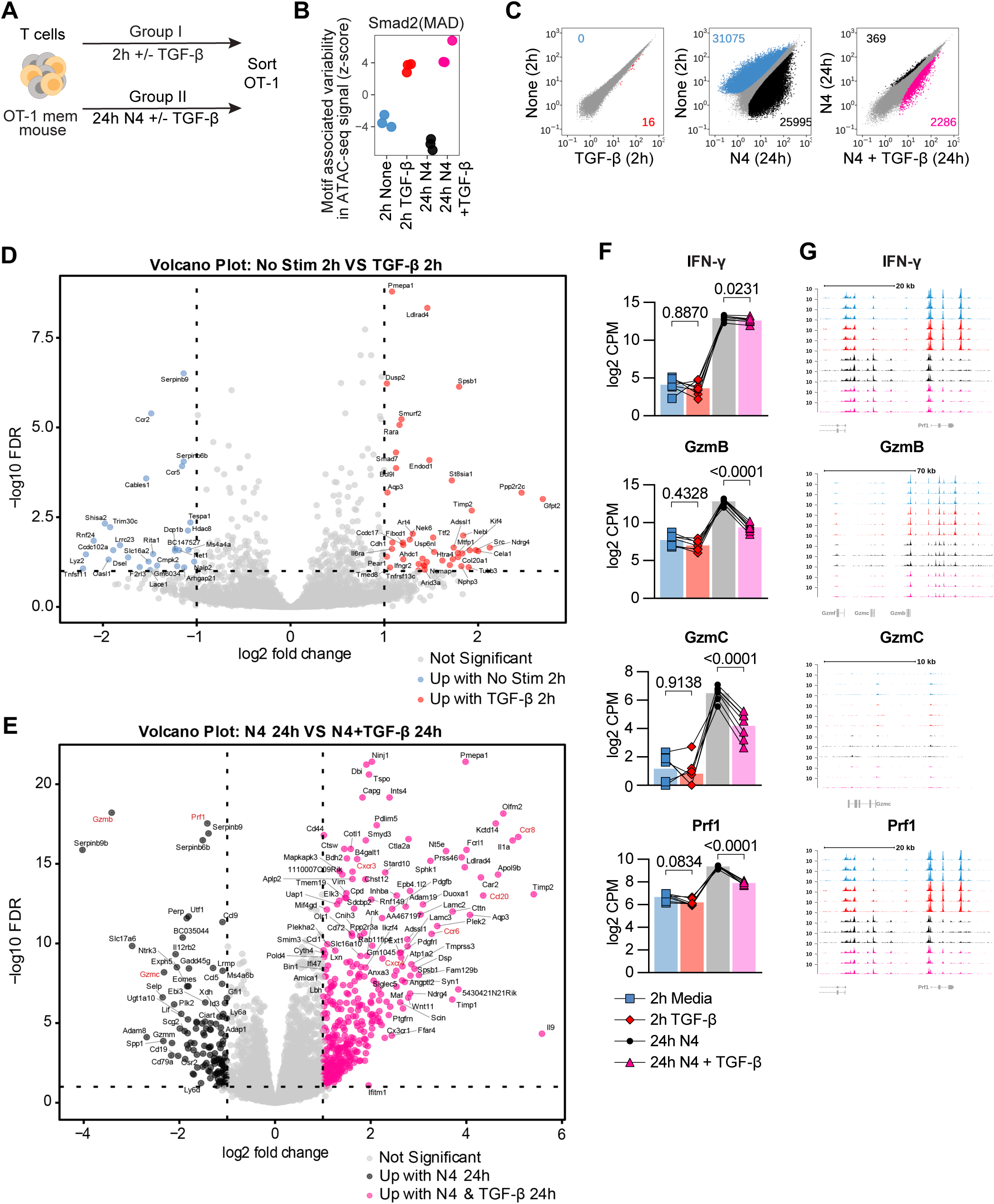
TGF-β epigenetically and transcriptionally alters memory CD8 T cell function. **(A)** Schematic for T cell isolation with magnet-activated cell sorting (MACS) from Ag-experienced OT-I memory mice, subsequent ex vivo stimulation, and sorting of Live, CD8+, CD45.1+ OT-I T_mem_ cells. Sorted OT-I T_mem_ were processed immediately for ATAC- and RNA-seq library preparation and sequencing. Stimulations were 2 hours with media alone or TGF-β at 100ng/mL and 24 hours 100nM N4 with or without TGF-β at 100ng/mL. **(B)** ChromVAR analysis of ATAC-seq signal (Z-score) at region containing the selected transcription factor motifs. **(C)** Scatterplot comparing differentially accessible chromatin regions for pairs of stimulation conditions. **(D)** Volcano plot depicting differentially expressed genes between 2 hour media and **(E)** 24 hour 100ng/mL N4 stimulation conditions: with TGF-β or without TGF-β. DE gene cutoff values were: adj p value 0.1, Log FC >1 and <-1. **(F)** Selected DE genes from RNAseq and indicated statistical significance. **(G)** Chromatin accessibility of selected genes from ATAC-seq. All data depicted are from n = 7 animals and 2 independent experiments.

### TGF-β alters the chemotactic properties of memory CD8 T cells

Several chemokines and chemokine receptors were also altered by TGF-β, including increased transcript expression of CCR8, CXCR3, CCR6, CXCR4 and CCL20. Of note, we detected a change in chromatin accessibility for CCR8, CXCR3, CCR6, CXCR4 and CCL20 indicating that the TGF-β-induced differences in chemokine transcripts may be due to increased access of their loci (**Figure 5A and 5B)**. Next, we examined if these alterations also resulted in changed protein expression in a set of follow up experiments. We performed ex vivo stimulations on OT-I T_mem_ as described in **Figure 3A** and **4A**. We found a modest increase of CXCR3 when OT-I T_mem_ were reactivated via their TCR and in the presence of TGF-β (**Suppl Fig 6A)**. In contrast, the changes for CCR8 were much more pronounced: we found that CCR8 had distinct low and high expression patterns dependent on the stimulation condition, and we gated these populations accordingly (**Suppl Fig 9B**). In context of TCR-mediated reactivation, TGF-β greatly increased the frequency of CCR8hi expressing OT-I T_mem_, but was most pronounced in the N4 and Q4-reactivated groups (**Figure 5C)**. Of note, this occurred even when TGF-β was added 6 or 12 hours after the reactivation stimulus (**Figure 5D)**. In contrast, TGF-β only elicited a substantial CCR8hi expressing OT-I T_mem_ population when given concurrently with the cytokines (**Figure 5D)**. Similarly, pre-exposure of OT-I T_mem_ to TGF-β for 2 hrs followed by washing out the TGF-β and reactivation with N4 or Q4 was sufficient to induce CCR8 expression that was nearly indistinguishable from the positive control groups (**Suppl Figure 6B and 6C)**. We also measured CCR8 expression in context of a TGF-β titration and found that the weaker the TCR activating signal, the higher CCR8hi expression frequency among OT-I T_mem_ (**Suppl Figure 6D)**. Thus, the CCR8 expression pattern is a negative mirror of the granzyme B expression data. Overall, these data highlight that TGF-β can modify the chemotactic properties of reactivated CD8 T_mem_ in a dose-dependent and reactivation signal-dependent manner.

**Figure 5.**
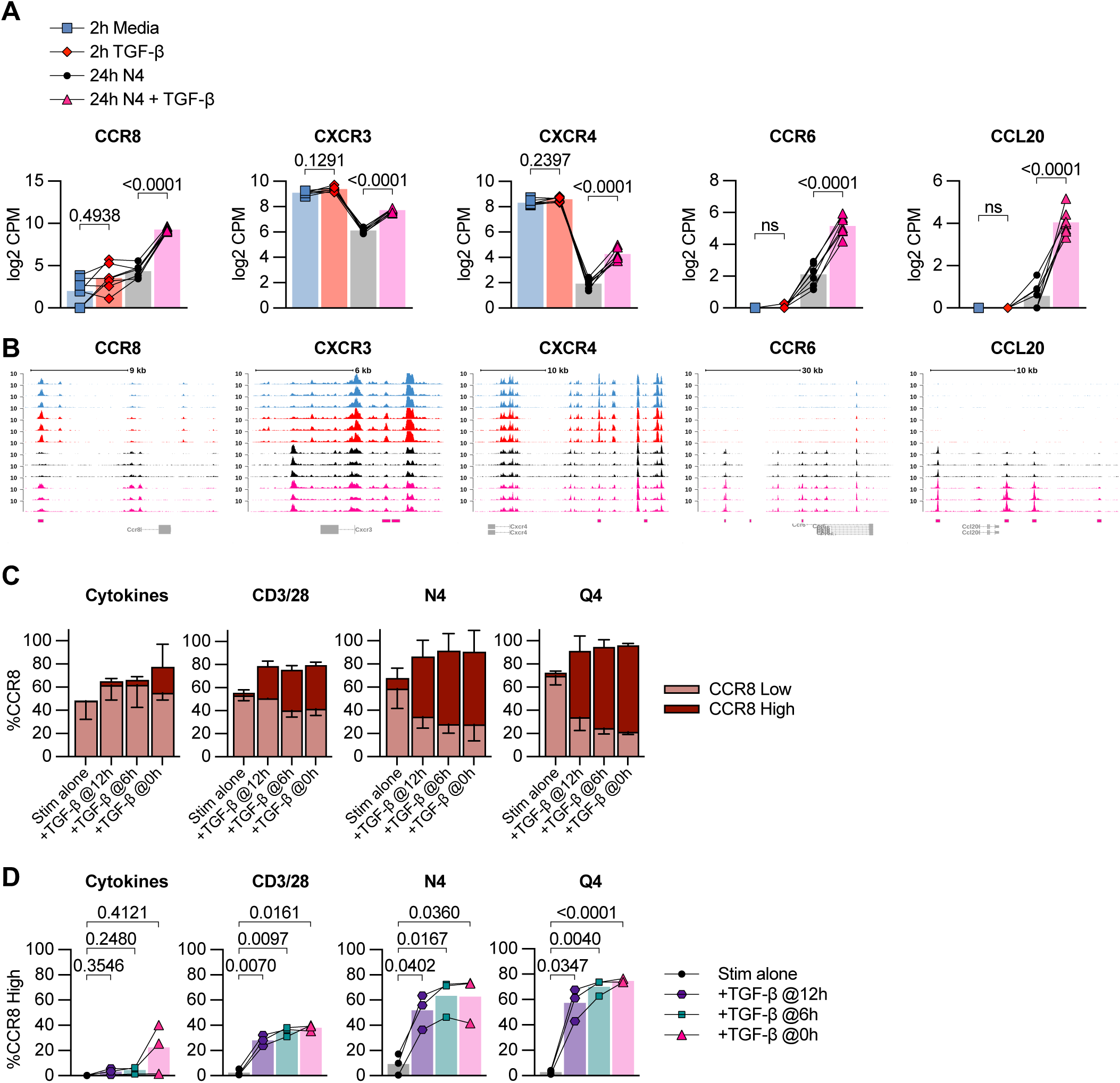
TGF-β epigenetically and transcriptionally alters memory CD8 T cell chemotaxis. **(A)** Selected DE genes from RNA-seq and indicated statistical significance. **(B)** Chromatin accessibility of selected genes from ATAC-seq. **(C)** Frequency of low and high CCR8 expression by flow cytometry in OT-I T_mem_ across 24h stimulation conditions with TGF-β addition at indicated timepoints. **(D)** Frequency of CCR8-high expression plotted individually and indicated statistical significance. Stimulations in **(C and D)** are from the same experiments shown in Fig. 3A. Indicated statistical significances were calculated using one-way Anova with Tukey’s multiple comparison test. In (**A and B**) RNA- and ATAC-seq data depicted are from n = 7 animals. In **(C)** and **(D)** data depicted are from n = 3. Data shown are from 2 independent experiments.

## Discussion

Genetic ablation approaches of TGF-β receptor signaling have provided a set of important tools to demonstrate that TGF-β signals directly act on T cells during priming and control survival, differentiation, effector function and formation of tissue-resident T cells (5, 6, 8-11, 25). A study by Ma and Zhang demonstrated that TGF-β signals are necessary for the proper maintenance of functional memory T cells (11), which has recently also been extended to chronic infections (26, 27). Importantly, dose-dependent effects of a ligand cannot be assessed with these genetic ablations models. TGF-β availability changes during inflammatory processes (28), but how these changes impact memory T cell function is poorly understood. We thus wanted to assess how low to high concentrations of TGF-β affect CD8 T_mem_ function in context of different reactivation signals. Reactivation of memory CD8 T cells is a critical component of providing protection against infections (29, 30), PD-1/PD-L1 induced anti-tumor responses (18) as well as vaccines targeting cancer (16). TGF-β has been reported to inhibit Ca2+ influx (31) thus indicating that the reactivation signal itself may affect the consequences of TGF-β signaling. The quality of the TCR signal controls the downstream transcriptional changes (32) and we considered that CD8 T_mem_ can be reactivated by a range of different TCR- as well as cytokine-mediated signals. To simultaneously manipulate reactivation and TGF-β signals, we needed to generate CD8 T_mem_ with intact TGF-β signaling and then control TGF-β and reactivation signals in an ex vivo set up.

When we reactivated OT-I T_mem_ by TCR cross-linking by plate bound antibodies, TGF-β very effectively inhibited granzyme B expression (1,3 ng/ml were sufficient for a 2-fold reduction in the frequency of granzyme B+ OT-I T_mem_). In contrast, when we reactivated OT-I T_mem_ with either N4 or Q4 peptide, we found that a 10-fold lower concentration of TGF-β was already sufficient to reduce the frequency of granzyme B+ OT-I T_mem_ 2-fold (0.16 and 0.09 ng/ml of TGF-β for N4 and Q4, respectively). These data strongly suggest that lower affinity responders are particularly susceptible to losing cytotoxic function, which is an important consideration for anti-tumor responses. This potent suppression of cytotoxic function of low affinity CD8 T cells then also begs the question how low affinity T cells could possibly contribute to pathogen clearance. A previous study indicated a potential decrease in total TGF-β in blood in the first days following infection with *Listeria monocytogenes* (LM) (28) and we similarly observed a decrease of total TGF-β in the spleen following infection with LM from 10.1 ng/g tissue during homeostasis, and decreased to 4.5 ng/g tissue 3 days following infection with Listeria monocytogenes (**Supplemental Figure 1A**). Of note, active TGF-β is often only a fraction of total TGF-β (13). Such an infection-associated decrease in TGF-β may be critical to allow for low affinity CD8 T_mem_ to exert cytotoxic function. It is also worthwhile to consider that such a decrease in active TGF-β represents a window of opportunity for self reactive T cells to acquire cytotoxic function. An association of viral infection and an autoimmune response was first suggested 40 years ago with autoreactive antibodies (33), but has since been demonstrated for T cells as well (34). This is typically thought to be the result of molecular mimicry between viral and self antigen, which could be facilitated during a decline in active TGF-β availability (10). Since infections also elicit cytokine-driven activation of CD8 T_mem_ (35-37), we examined how bystander-activated CD8 T_mem_ are affected by TGF-β signals. Interestingly, TGF-β had a similar effect on CD8 T_mem_ reactivated with IL-12, 15 and 18: the reduction in gzmB expression was comparable to CD3/CD28 cross-linking (1,3 ng/ml were sufficient for a 2x reduction in the frequency of granzyme B+ T cells) with a high concentration of pro-inflammatory cytokines, but the susceptibility to TGF-β-mediated inhibition of cytotoxicity increased as we decreased cytokine concentrations. Overall, these data highlight the importance of the strength of the activating signal in regards to the ability of TGF-β to inhibit cytotoxic function.

Based on studies that relied on priming on naïve T cells or used T cell clones, it is often assumed that TGF-β concurrently inhibits IFN-γ and cytotoxic function (25). However, across all experimental conditions, we consistently observed that the impact of TGF-β signals on IFN-γ expression by reactivated CD8 T_mem_ was rather limited. This distinct effect of TGF-β on granzyme B and IFN-γ expression in reactivated CD8 T_mem_ is curious, particularly in context of the tumor microenvironment with presumably abundant active TGF-β. Our data indicate that TGF-β can inhibit direct cytotoxicity by reactivated CD8 T_mem_, but IFN-γ could still allow for myeloid cell-mediated tumor killing (38, 39). In context of an infection, this selective disabling of cytotoxicity could limit pathology while still allowing for IFN-γ mediated protective effects and continued recruitment of immune cells (40).

In our initial set of experiments, we provided reactivation and TGF-β signals at the same time, but we considered that CD8 T_mem_ may receive activating signals before or after a TGF-β signal (for example, reactivation in a lymph node followed by TGF-β exposure in the tissue, or vice versa). Since our ex vivo experimental system allowed us to have temporal control of the sequence of signaling events (TGF-β exposure before, together with or after the activation event), we explored these different scenarios. We found that receiving TGF-β signals after reactivation still efficiently reduced cytotoxicity and, similarly, brief exposure to TGF-β prior to an activation event was sufficient to reduce cytotoxic effector function. In our system this suppressive effect lasts for 24 hours, but this observation of course begs the question of how long the decrease in cytotoxic function may last in vivo. Defining the duration of suppression will be important in follow up studies and is relevant in context of the association between viral infection and autoimmune responses, as well as anti-tumor responses. Based on these data, we speculated that TGF-β may alter chromatin accessibility.

We did not observe changes in chromatin accessibility to perforin or granzyme genes, which were significantly decreased in abundance in the presence of TGF-β, but interestingly detected epigenetic changes for several chemokine receptors, including CCR6, CXCR3, CXCR4 and CCR8. These data suggest that at least some of the TGF-β-mediated changes are epigenetic in nature. We observed a TGF-β-mediated increase in CXCR3 expression in context of CD8 T T_mem_ reactivation, while a recent study reported that deletion of TGF-βRI driven by CD8a-cre enhanced CXCR3 expression on CD8 T cells(41). A possible explanation for this difference is due the timing of deletion as noted in other TGF-β studies in regards to T cell activation and differentiation (10, 11). We were particularly interested in CCR8 expression, which has often been observed on intratumoral regulatory T cells (42). Thus, TGF-β could potentially push reactivated CD8 T_mem_ to co-localize with these Tregs in tumors thereby ensuring continued control over their effector function. It could also be a critical signal to route CD8 T_mem_ to the skin, which is a physiological target site for CCR8+ T cells (43). Of note, induction of CCR8 expression was TGF-β dose-dependent and even low doses of 0.04-0.06 ng/ml were sufficient to elicit expression in about 50% of OT-I T cells reactivated with N4 or Q4, respectively. TGF-β also increased expression of the adhesion receptor ninjurin-1 (Ninj1), which is involved in T cell crawling in blood vessels (44), metalloproteinase 1 (Timp1) and the metalloprotease Meltrin β (ADAM19) and the chemokine CCL20, which orchestrates interactions with CCR6-expressing immune cells subsets (including Tregs, Th17 and dendritic cells)(45, 46). In addition to gene expression changes related to cell motility and trafficking, GO analysis also revealed changes related to cell metabolism (**Suppl Fig 5C**).

We were surprised by the large number of regulatory elements that changed during reactivation (almost 26000). About 10% of these elements were affected by TGF-β 24hrs after reactivation, indicating that TGF-β signals are not merely a specific suppressor of effector function, but rather a modifier CD8 T_mem_ function, which is highly relevant in regards to blocking TGF-β signaling for therapeutic purposes. Targeting TGF-β for therapeutic purposes, specifically to alter immune responses, is of great clinical interest, but the pleiotropic properties of TGF-β across different cell types have complicated these efforts (1, 47). Advances in the design of biologic therapeutics now allow for a more specific targeting of cells to block or activate receptor function (47), but our data highlight that that even for CD8 T_mem_ inhibition of TGF-β signals does not simply equal increased functionality: for example, complete blocking of TGF-β may preclude CCR8 expression and prevent trafficking to sites in which ligands (including CCL1 and CCL8) are expressed (42, 48). This includes trafficking to the skin, which would presumably interfere with targeting melanomas (49), but also trafficking to the CCL8^+^ hypoxic regions of solid tumors (50).

Overall, our data indicate that TGF-β should not be considered a suppressor of effector function for CD8 T_mem_, but rather a modifier of CD8 T_mem_ function in the context of reactivation. Our data support the notion that TGF-β does not affect all CD8 T_mem_ equally since the functional consequences of a TGF-β signal are shaped by the strength of the reactivation signal. Finally, our data also highlight that TGF-β signals can exert their function regardless if they are received before or after the reactivating event, which is an important consideration for interpreting studies that assess CD8 T_mem_ function in situ.

## Methods

### Mice

Mouse protocols and experimentation conducted at the Fred Hutchinson Cancer Research Center were approved by and in compliance with the ethical regulations of the Fred Hutchinson Cancer Research Center’s Institutional Animal Care and Use Committee. All animals were maintained in specific pathogen-free facilities and infected in modified pathogen-free facilities. Experimental groups were non-blinded and animals were randomly assigned to experimental groups. We purchased 6-week-old female C67BL/6J mice from the Jackson Laboratory; OT-I mice were maintained on CD45.1 congenic backgrounds. To generate OT-I memory mice, we adoptively transferred 1 × 10^4^ OT-I T cells in sterile 1× PBS i.v. per C57BL/6J recipient, and subsequently infected recipients i.v. with 1 × 10^6^ PFU OVA-expressing vesicular stomatitis virus (VSV-OVA) (51) or 4 × 10^3^ CFU OVA-expressing *Listeria monocytogenes* (LM-OVA) as previously described (52). We allowed ≥ 60 days to pass after initial VSV or ≥ 30 days LM infections before assaying tissues.

### T cell isolation and ex vivo stimulation

To enrich bulk T cells from single cell suspensions, we used mouse-specific and human-specific T cell negative isolation MACS (STEMCELL Technologies, Canada). We plated 0.5–1 × 10^6^ T cells per well in 96-well V-bottom tissue culture plates. We cultured cells in mouse RP10 media (RPMI 1640 supplemented with 10% FBS, 2mM L-glutamine, 100 U/mL penicillin-streptomycin, 1mM sodium pyruvate, 0.05mM β-mercaptoethanol, and 1mM HEPES) or human RP10 (RPMI 1640 supplemented with 10% FBS2mM L-glutamine, and 100 U/mL penicillin-streptomycin). To stimulate cells, we cultured mouse T cells in mouse RP10 with recombinant mouse TGF-β1 (Biolegend Cat # 763104), rIL-12, rIL-15, and rIL-18 (BioLegend) (at specified concentrations), with plate-bound anti-CD3 and anti-CD28 antibodies (prepared by incubating plates for 2 hours at 37°C with antibodies at 500ng/mL and 1ug/mL respectively), with N4, with Q4, or with media alone. For human T cell stimulations, we used human RP10 media with recombinant human TGF-β1 (PeproTech Cat #100-21), Dynabeads human T-Activator (Thermo Fisher) anti-CD3/CD28 beads (at a 1:1 bead/cell ratio), or with media alone. We cultured cells at 37°C, 5% CO2, sampling cells at 0, 6-8, and 24 hours for flow staining. To measure secreted cytotoxic molecules, we stimulated T cells in the presence of GolgiPlug (BD Biosciences) (1:1000 dilution) for the final 4 hours of stimulation, after which we conducted intracellular cytokine staining (ICS).

### Flow cytometry

We conducted all flow staining for mouse and human T cells on ice and at room temperature, respectively. All mouse and human flow panel reagent information, stain conditions, and gating are included in **(Supplemental Fig. 7, 8, 9 Supplemental tables 1,2,3)**. We conducted LIVE/DEAD fixable aqua (AViD) staining in 1× PBS. For surface staining, we utilized FACSWash (1 × PBS supplemented with 2% FBS and 0.2% sodium azide) as the stain diluent. We fixed cells with the FOXP3 fixation/permeabilization buffer kit (Thermo Fisher) and conducted intranuclear stains using the FOXP3 permeabilization buffer (Thermo Fisher) as diluent. For ICS panels, we fixed cells with Cytofix/Cytoperm (BD Biosciences) and conducted intracellular stains using Perm/Wash buffer (BD Biosciences) as diluent. We resuspended cells in FACSWash and acquired events on a FACSSymphony, which we analyzed using FlowJo v10 (BD Biosciences). We conducted statistical testing using Prism v8 (GraphPad).

### ELISA for TGF-β

Female C57BL/6J mice were infected with 4 × 10^3^ CFU LM-OVA. Spleens were weighed then mechanically dissociated in 500uL buffer (1x PBS supplemented with 0.05% Tween) with scissors in a microcentrifuge tube. To separate debris, samples were centrifuged at 1000 g for 10 minutes at 4°C and the supernatants were stored at -80°C until assay. Total TGF-β levels were determined by acid activation of the latent TGF-β1 in the sample using the sample activation kit 1 (DY010) (R&D Systems).

### Bulk RNA Sequencing

Bulk RNA-seq was performed on 500 sort-purified OT-I T cells derived from OT-I memory mice after culture in conditions of 2 hours no stimulation, 2 hours stimulation with 100ng/mL TGF-β, 24 hours stimulation with 100nM N4, and 24 hours stimulation with 100nM N4 and 100ng/mL TGF-β. 24 hour stimulation stain control was performed to ensure T cell activation occurred consistently with prior experiments (**Suppl Fig 7A**). In total, 28 samples were sequenced, and each condition was represented by a total of 7 biological replicates (combined from 2 independent experiments). Cells were prepared for RNA sequencing and data were overall analyzed as previously described (53) aside from using the GRCm38 reference genome.

### ATAC Sequencing

ATAC-seq was performed on pools of 40,000 to 50,000 sort-purified OT-I T cells pooled from 2 to 3 mice. DNA was purified as previously described (54). Fastq files were used to map to the mm10 genome using the ENCODE ATAC-seq pipeline (55), with default parameters, except bam files used for peak calling were randomly downsampled to a maximum of 50 million mapped reads. Peaks with a MACS2 (56) computed q value of less than 0.0001 in at least one replicate were merged with bedtools (57) function intersect and processed to uniform peaks with the functions getPeaks and resize from R package ChromVAR (58). Reads overlapping peaks were enumerated with getCounts function from ChromVAR and normalized and log2-transformed with *voom* from R package *limma (59, 60)*. Peaks with 6 or more normalized counts per million mapped reads at least one replicate were included to define a global peak set of 99,317 peaks. Pairwise Euclidean distances were computed between all samples using log2-transformed counts per million mapped reads among the global peak set. Differentially accessible peaks were identified in pairwise comparisons based on fdr adjusted p values of less than 0.05, fold change of at least 1.5 and with an average of 6 normalized counts per million mapped reads using R package *limma.* Motif associated variability in ATAC-seq signal was computed with R package ChromVAR. Genome-wide visualization of ATAC-seq coverage was computed with deeptools (61) function coveragebam, using manually computed scale factors based on the number of reads within the total peak set.

## Acknowledgements

We would like to thank members of the Prlic lab for critical discussion. We thank the Flow Cytometry Shared Resources of the FHCRC and the Genomics Core Lab of the Benaroya Research Institute for sequencing. This work was supported by NIH grants R01AI123323 (to M.P.) and R01AI151021 to (J.S.-B.). The experimental overview schemes in Fig 1A, 4A, and supplemental Fig. 2A use templates from BioRender.com.

## Author contributions

AT designed and performed experiments, analyzed data and co-wrote the manuscript. AK designed and performed experiments, analyzed data and edited the manuscript. JSB analyzed data and co-wrote the manuscript.

MP designed the study, analyzed data and co-wrote the manuscript.

