## Supplemental Figures for "TGF-β broadly modifies rather than specifically suppresses reactivated memory CD8 T cells in a dose-dependent manner"

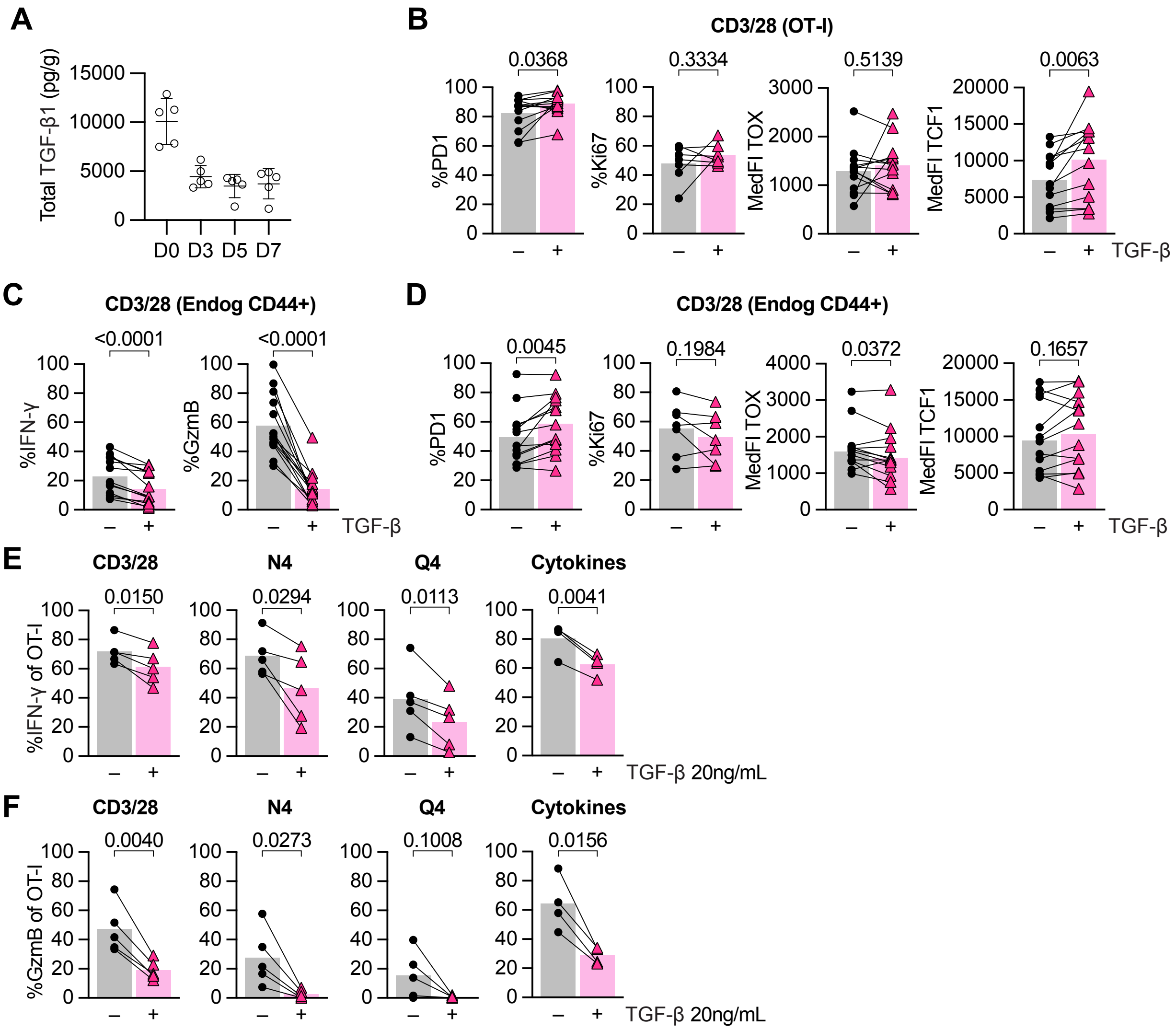


**Supplemental Figure 1: TGF-β increases PD-1 but not other markers of activation on memory CD8+ T cells**

See Figure 1A for experimental setup. Enriched T cells were stimulated for 24 hours with CD3/28 in the presence or absence of TGF-β at 100ng/mL. **(A)** Total levels of extracellular TGF-β1 measured by ELISA from spleen of C67BL/6J mice at indicated infection timepoints (n = 5 animals). Infection was 4 × $10^3$ CFU LM-OVA on Day 0. **(B)** Frequencies of PD1 and Ki67 and median fluorescence intensity (MedFI) of Tox and TCF1 in OT-1 Tmem. **(C)** Frequencies of IFN-γ, GzmB, **(D)** PD1, Ki67, and MedFI of Tox and TCF1 in endogenous CD44+ T cells. In **(B - D)** n = 13 animals for all markers except Ki67 n = 7. **(E)** Frequencies of IFN-γ and **(F)** GzmB of OT-I Tmem at 24h across indicated stimulation conditions with or without TGF-β at 20ng/mL. In **(E and F)** CD3/28, N4, and Q4 n = 5 animals, and Cyt n = 4. Statistical significances were calculated using paired t tests. Data shown are from 4 to 11 independent experiments.

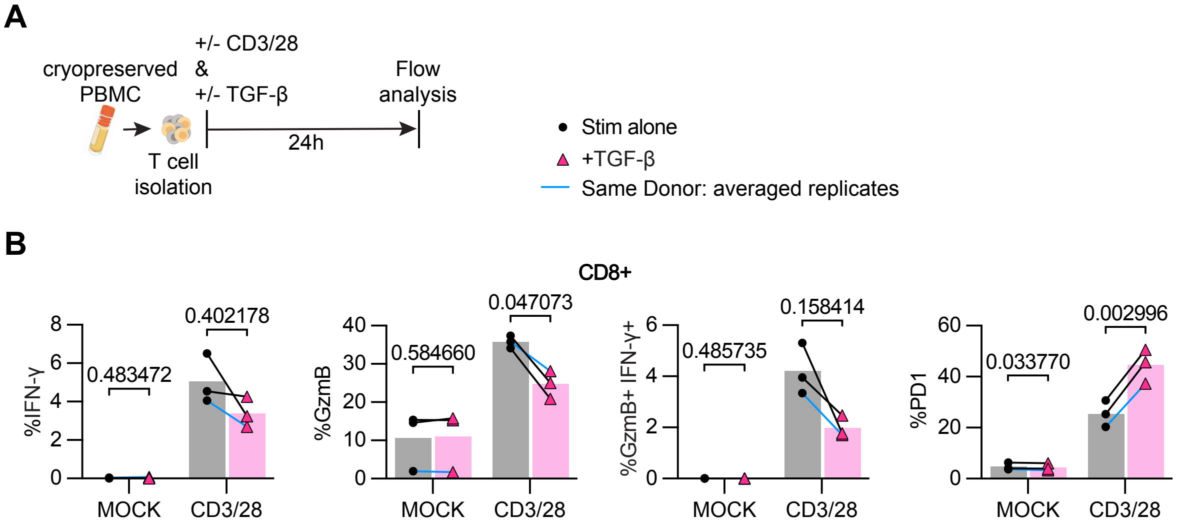


**Supplemental Figure 2: TGF-β inhibits cytotoxicity of CD8+ T cells from human PBMC**
**(A)** Schematic of T cell isolation with magnet-activated cell sorting (MACS) from human PBMC and subsequent ex vivo stimulation and analysis. Stimulation was 24 hours with anti-CD3/CD28 microbeads in the presence or absence of TGF-β at 100ng/mL. **(B)** Frequencies of indicated markers in all CD8+ T cells (n = 3 donors). Experimental repeats belonging to the same donor are averaged and indicated by the blue (4 replicates). Statistical significances were calculated using multiple paired t tests. Data shown are from 4 independent experiments.

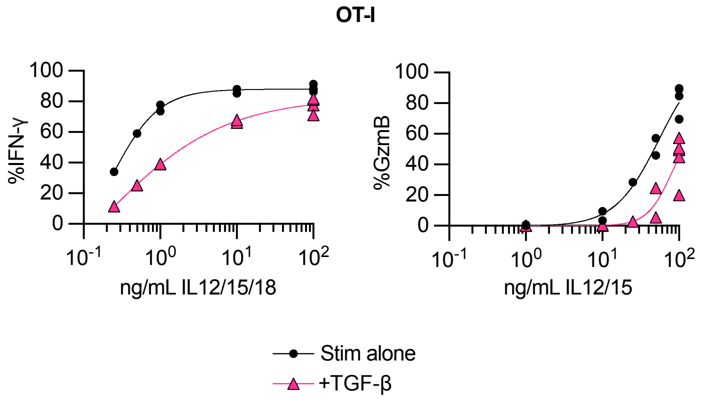


**Supplemental Figure 3: Pro-inflammatory cytokine abundance determines the susceptibility of OT-I Tmem to inhibition by TGF-β**

See figure 1A for experimental setup. Stimulation was 24h with IL-12, IL-15, and IL-18 or IL-12 and IL-15 at the indicated (titrated) concentrations with or without TGF-β at 100ng/mL. Frequencies of IFN-γ and GzmB by OT-I $T_{mem}$ (n = 3 animals). Data shown are from 3 independent experiments.

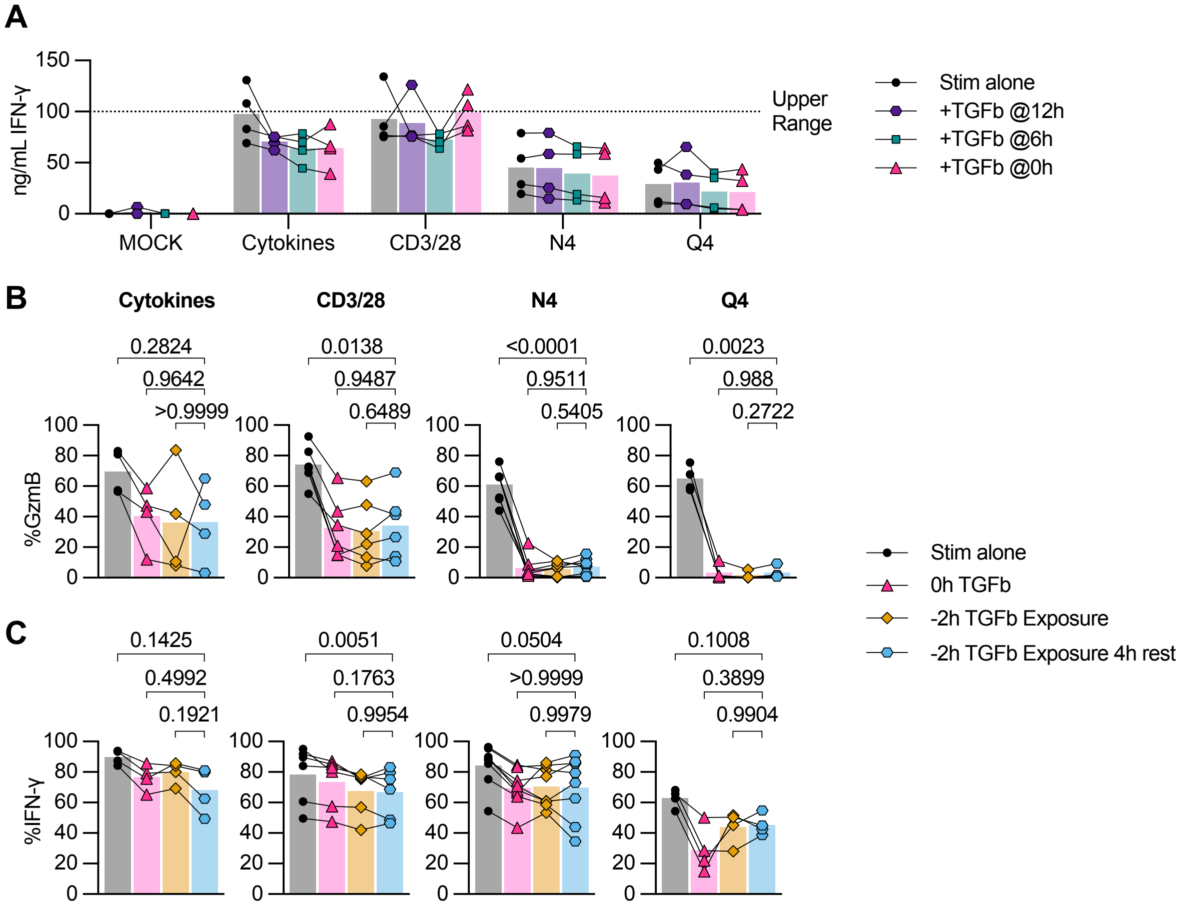


**Supplemental Figure 4: TGF-β exposed OT-I Tmem do not regain full cytotoxic function after short rest period**

**(A)** IFN-γ levels measured by ELISA from supernatant of ex vivo experiments described in Figure 3A (n = 4 animals). **(B and C)** Stimulated OT-I $T_{mem}$ from experiments outlined in Figure 3D including an additional condition of a 4-hour rest in fresh media between TGF-β exposure and activating stimulation. **(B)** Frequencies of GzmB and **(C)** IFN-γ in OT-I $T_{mem}$ compared across stimulation conditions (CD3/28 n = 6 animals, N4 n = 8, Q4 and Cyt n = 4). All indicated statistical significances were calculated using one-way ANOVA. Data shown are from 3 to 7 independent experiments.

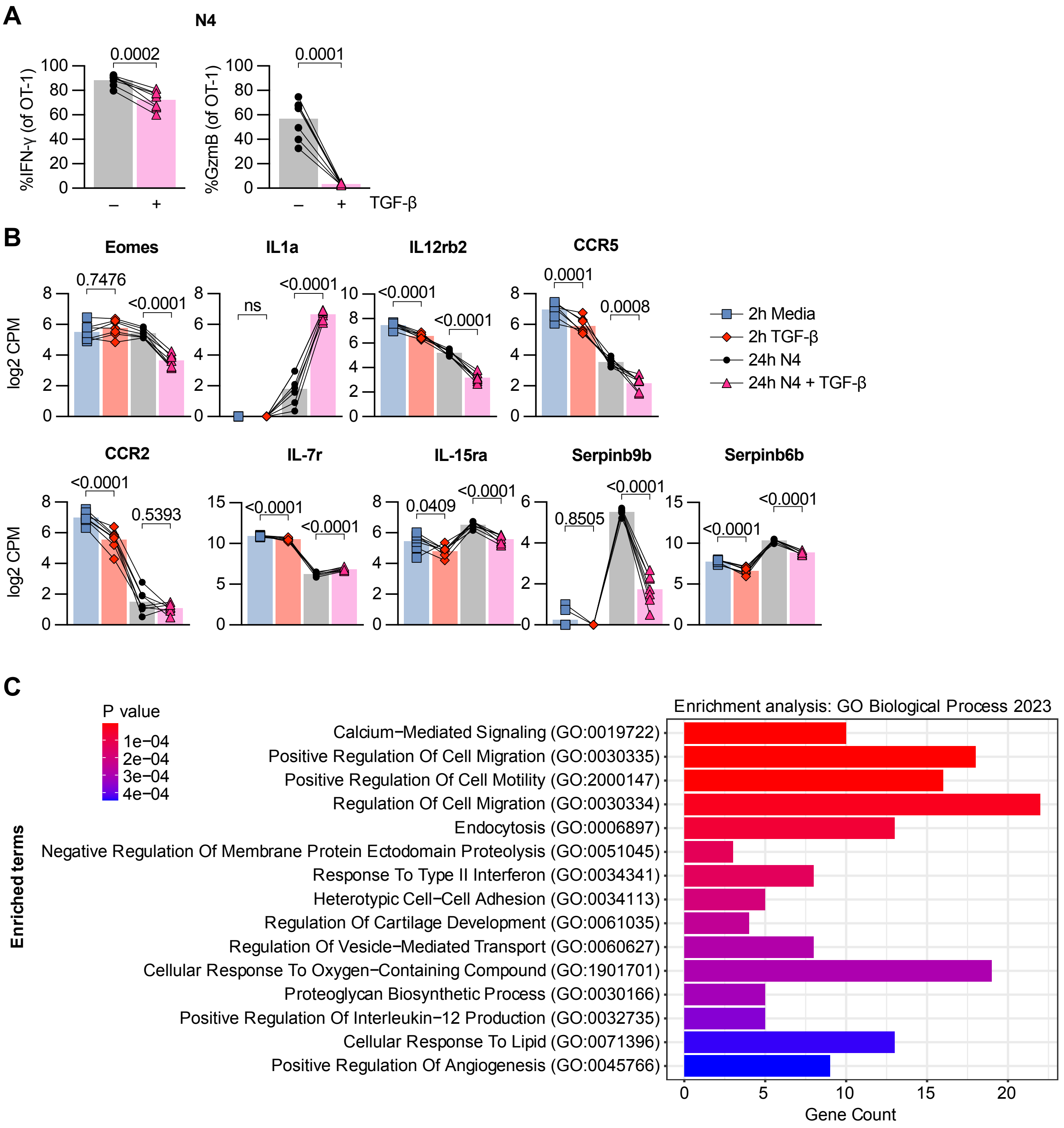


**Supplemental Figure 5: TGF-β induces transcriptional changes associated with cell migration in OT-I Tmem**

**(A)** Frequencies of IFN-γ and GzmB in OT-I Tmem from mice used for ATAC- and RNA-seq after 24h N4 stimulation with or without TGF-β at 100ng/mL (n = 7 animals). Statistical significances were calculated using paired t tests. **(B)** Selected DE genes from RNA-seq and calculated adj. P values. **(C)** Enrichment analysis of RNAseq data from GO Biological Process 2023 database. Data shown are from 2 independent experiments.

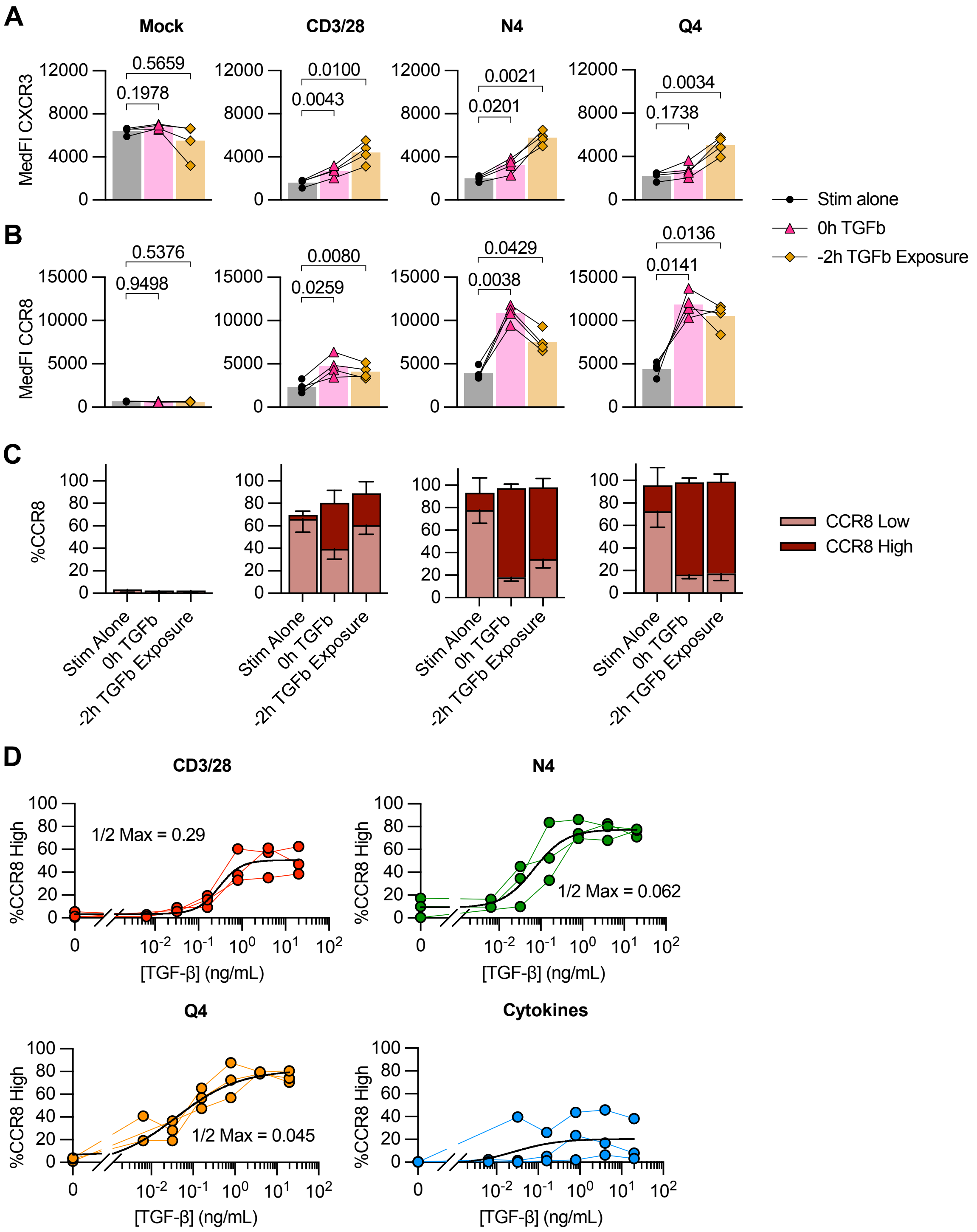


**Supplemental Figure 6: TGF-β induces chemokine receptor expression in OT-I Tmem in a dose-dependent manner**

**(A - C)** See Figure 3D for experimental setup. **(A)** MedFI CXCR3 and **(B)** MedFI CCR8 in OT-1 $T_{mem}$. **(C)** Frequency of low and high CCR8 expression by flow cytometry in OT-I $T_{mem}$ (n = 4 animals). **(D)** Frequency of CCR8-high expression by flow cytometry across stimulation conditions in the presence of titrated TGF-β (n = 3). TGF-β was titrated in five-fold dilutions starting with 20ng/mL and ending at 0.032ng/mL. Data shown are from 2 to 3 independent experiments.

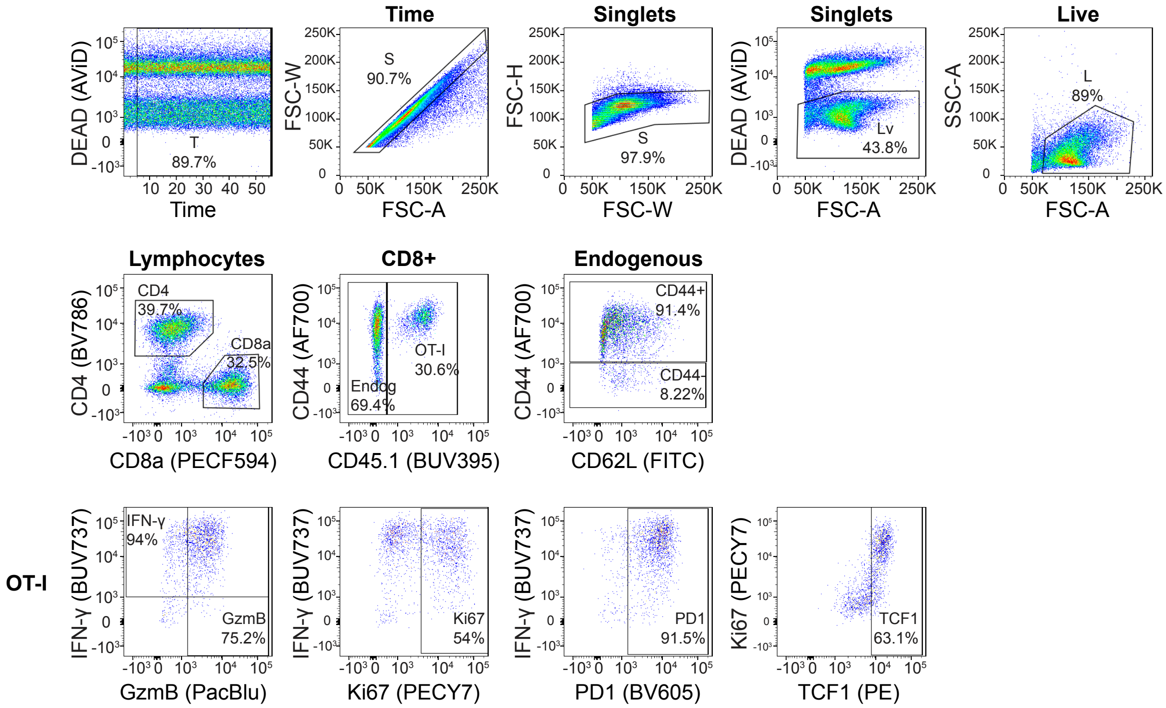


**Supplemental Figure 7: Representative flow gating for T cells isolated from OT-I memory mice**

Related to Fig.1, 2, 3, Supplemental Fig. 1, 3, 4. Stimulation was 24h with plate-bound CD3/28.

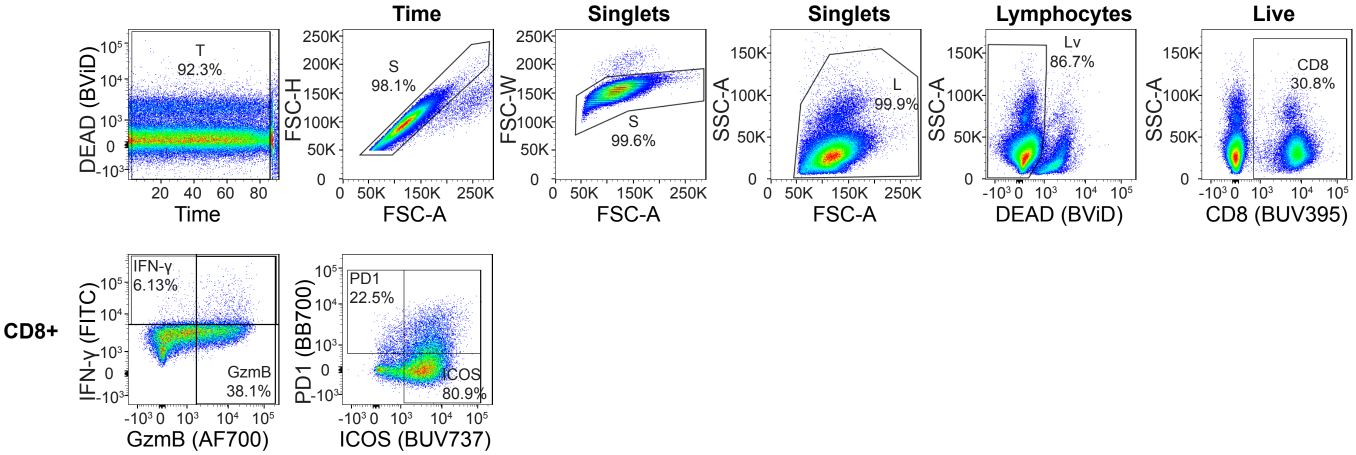


**Supplemental Figure 8: Representative flow gating for T cells isolated from human PBMC**
Related to Supplemental Fig. 2. Stimulation was 24 hours with anti-CD3/CD28 microbeads.

A

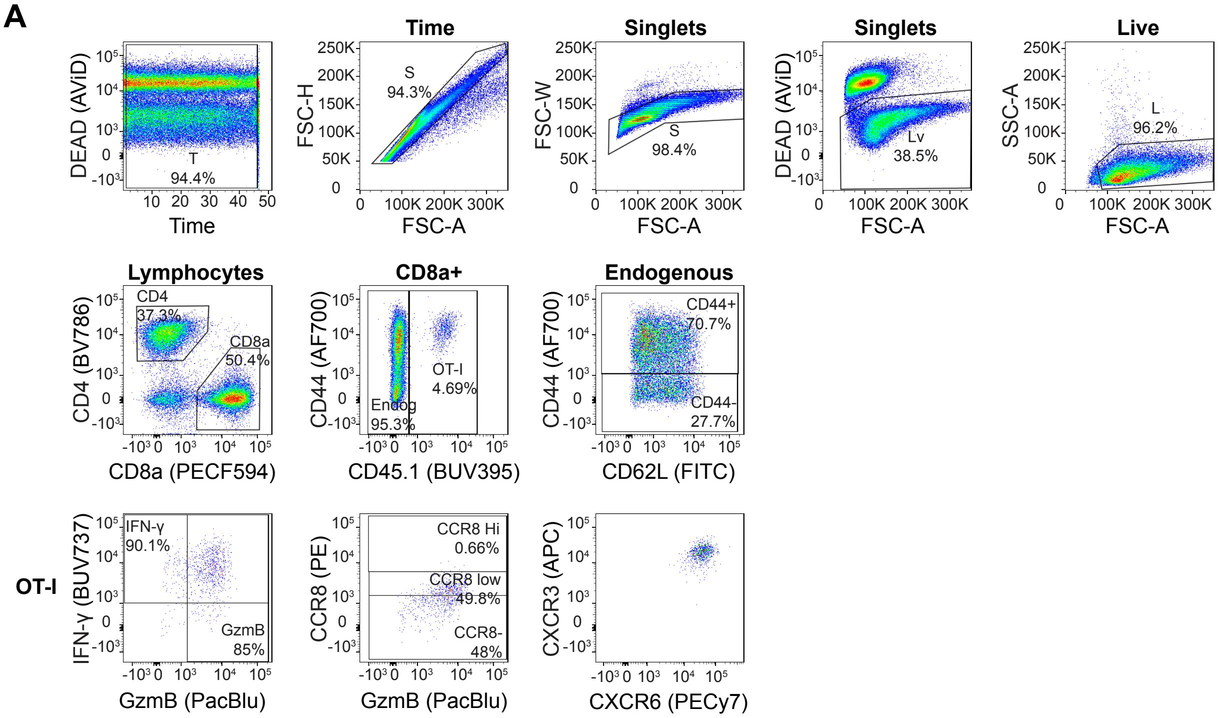


B

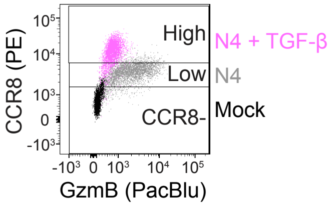


**Supplemental Figure 9: Representative flow gating for T cells isolated from OT-I memory mice**

Related to Fig. 5, Supplemental Fig. 6. **(A)** Stimulation was 24h with plate-bound CD3/28. **(B)** Representative gating of CCR8 expression in OT-I Tmem across stimulation conditions: Media alone (Mock), N4, and N4 + TGF-β.

| TGF-β Mouse Panel | | | | | |
|---|---|---|---|---|---|
| Reagent | Fluor | Dilution | Clone | Vendor | Cat.no |
| Viability stain (PBS diluent) 20 min, on ice | | | | | |
| LIVE/DEAD fixable aqua viability dye (AViD) | V510 | 1:250 | NA | Thermo Fisher | Cat#L34966 |
| Surface stain (FACS Wash diluent) 20 min, on ice | | | | | |
| FC block (CD16/32) | Purified | 1:200 | 2.4G2 | BD Biosciences | Cat# 553141 |
| CD4 | BV786 | 1:200 | GK1.5 | BD Biosciences | Cat#563331 |
| CD8a | PECF594 | 1:300 | 53-6.7 | BD Biosciences | Cat#562283 |
| CD45.1 | BUV395 | 1:200 | A20 | BD Biosciences | Cat#565212 |
| CD44 | AF700 | 1:200 | IM7 | Thermo FIsher | Cat#560567 |
| PD-1 | BV605 | 1:100 | 29F.1A12 | Biolegend | Cat#135220 |
| CD62L | FITC | 1:200 | MEL-14 | eBioscience | Cat#11-0621-85 |
| Fix (eBioscience FOXP3 fixation buffer) 20 min, on ice | | | | | |
| Intracellular stain (eBioscience FOXP3 perm buffer diluent) 30 min, on ice | | | | | |
| Ki-67 | PECy7 | 1:800 | SolA15 | BioLegend | Cat#52426 |
| TOX | APC | 1:100 | REA473 | Miltenyi Biotec | Cat#130-118-335 |
| IFN-γ | BUV737 | 1:200 | XMG1.2 | BD Biosciences | Cat#612769 |
| GzmB | PacBlu | 1:100 | GB11 | BioLegend | Cat#515408 |
| TCF1/7 | PE | 1:40 | C63D9 | Cell Signaling | Cat#144565 |

**Supplemental Table 1: TGF-β mouse panel**
Related to Fig.1, 2, 3, Supplemental Fig. 1, 3, 4. Mouse flow cytometry panel for T cells isolated from OT-1 memory mice.

| TGF-β Human Panel | | | | | |
|---|---|---|---|---|---|
| Reagent | Fluor | Dilution | Clone | Vendor | Cat.no |
| Viability stain (PBS diluent) 20 min, room temp | | | | | |
| Human TruStain FcX (Fc-Block) | NA | 1:25 | NA | Biolegend | Cat#422302 |
| LIVE/DEAD fixable blue viability dye (BViD) | UV450 | 1:500 | NA | Thermo Fisher | Cat#L34962 |
| Surface stain (Brilliant Stain Buffer diluted 10x in FACS Wash) 20 min, room temp | | | | | |
| CD8 | BUV395 | 1:80 | RPA-T8 | BD Biosciences | Cat#563795 |
| CXCR6 | BUV563 | 1:20 | 13B 1 E5 | BD Biosciences | Cat#748450 |
| CCR7 | BUV661 | 1:80 | 2-L1-A | BD Biosciences | Cat#749824 |
| ICOS | BUV737 | 1:20 | DX29 | BD Biosciences | Cat#749665 |
| CD25 | BV421 | 1:40 | 2A3 | BD Biosciences | Cat#564033 |
| CD28 | BV480 | 1:40 | CD28.2 | BD Biosciences | Cat#566110 |
| CD45RA | BV570 | 1:160 | HI100 | BioLegend | Cat#304132 |
| CD39 | BV605 | 1:40 | A1 | BioLegend | Cat#328236 |
| CD69 | BV650 | 1:20 | FN50 | BD Biosciences | Cat#310933 |
| CD103 | BV750 | 1:160 | Ber-ACT8 | BD Biosciences | Cat#747099 |
| CCR5 | BV785 | 1:20 | 3A9 | BD Biosciences | Cat#565001 |
| PD1 | BB700 | 1:20 | EH12.1 | BD Biosciences | Cat#566460 |
| IL-1R1 | PE | 1:20 | polyclonal | R&D Systems | Cat#FAB269P |
| CXCR3 | PE-CF594 | 1:20 | 1C6/CXCR3 | BD Biosciences | Cat#562451 |
| CD137 | PE-Cy5 | 1:20 | 4B4-1 | BD Biosciences | Cat#551137 |
| IL18R1 | PE-Cy7 | 1:40 | H44 | BioLegend | Cat#313812 |
| Fix (eBioscience FOXP3 fixation buffer) 20 min, room temp | | | | | |
| Intracellular stain (eBioscience FOXP3 perm buffer diluent) 30 min, room temp | | | | | |
| IFN-γ | FITC | 1:40 | B27 | BD Biosciences | Cat#554700 |
| GzmB | AF700 | 1:80 | GB11 | BD Biosciences | Cat#560213 |
| CCR8 | BV711 | 1:20 | 433H | BD Biosciences | Cat#747575 |

**Supplemental Table 2: TGF-β human panel**
Related to Supplemental Fig. 2. Human flow cytometry panel for T cells isolated from human PBMC.

| Chemokine Receptor Panel: Mouse | | | | | |
|---|---|---|---|---|---|
| Reagent | Fluor | Dilution | Clone | Vendor | Cat.no |
| Viability stain (PBS diluent) 20 min, on ice | | | | | |
| LIVE/DEAD fixable aqua viability dye (AViD) | V510 | 1:250 | NA | Thermo Fisher | Cat#L34966 |
| Surface stain (FACS Wash diluent) 20 min, on ice | | | | | |
| FC block (CD16/32) | Purified | 1:200 | 2.4G2 | BD Biosciences | Cat#553141 |
| CD4 | BV786 | 1:200 | GK1.5 | BD Biosciences | Cat#563331 |
| CD8a | PECF594 | 1:300 | 53-6.7 | BD Biosciences | Cat#562283 |
| CD45.1 | BUV395 | 1:200 | A20 | BD Biosciences | Cat#565212 |
| CD44 | AF700 | 1:200 | IM7 | BD Biosciences | Cat#560567 |
| CD62L | FITC | 1:200 | MEL-14 | eBioscience | Cat#11-0621-85 |
| CXCR3 | APC | 1:100 | CXCR3-173 | BD Biosciences | Cat#562266 |
| CXCR6 | PECy7 | 1:200 | SA051D1 | BioLegend | Cat#151118 |
| Fix (eBioscience FOXP3 fixation buffer) 20 min, on ice | | | | | |
| Intracellular stain (eBioscience FOXP3 perm buffer diluent) 30 min, on ice | | | | | |
| CCR8 | PE | 1:200 | SA214G2 | Biolegend | Cat#150311 |
| IFN-γ | BUV737 | 1:200 | XMG1.2 | BD Biosciences | Cat#612769 |
| GzmB | PacBlu | 1:100 | GB11 | Biolegend | Cat#515408 |

**Supplemental Table 3: Chemokine receptor mouse panel**
Related to Fig. 5, Supplemental Fig. 6. Mouse flow cytometry panel for T cells isolated from OT-1 memory mice.
